# Genomic analysis of population structure, antimicrobial resistance, and virulent factors of methicillin-resistant *Macrococcus caseolyticus* in global lineages

**DOI:** 10.1101/2022.04.12.488041

**Authors:** Yu Zhang, Shengyi Min, Yuxuan Sun, Jiaquan Ye, Zhemin Zhou, Heng Li

**Author notes:** Corresponding author: Pasteurien College, Suzhou Medical College, Soochow University, Suzhou, Jiangsu, China, Email address.

## Abstract

*Macrococcus caseolyticus* is an opportunistic pathogen frequently detected in dairy products as well as veterinary infections. The present study examined the population structure, antimicrobial resistance, and virulent factors of methicillin-resistant *M. caseolyticus* isolates in retail meat from Shanghai (n=10) and global isolates from GenBank (n=87). All strains were divided into five lineages that distributed in Europe (82.4%, n = 80), Asia (11.3%, n = 11), North America (4.1%, n = 4), Oceania (1%, n = 1) and Africa (1%, n = 1). MLST typing revealed novel alleles in Chinese *M. caseolyticus* strains. Furthermore, a total of 24 AMR genes associated with 10 classes of antimicrobial agents were identified in the isolates from global lineages, carried by dominant plasmids such as rep7a, rep22 and repUS56. Comparing to other lineages, genomes from the Chinese lineage carried significantly more AMR genes (p<0.005) and less virulent factors (p<0.001), which may be explained by the local evolution of *M. caseolyticus* in China. Finally, scanning electron microscopy (SEM) and transmission electron microscopy (TEM) were enrolled for morphology comparison between *M. caseolyticus* and *S. aureus*, showing that *M. caseolyticus* has a larger diameter and thicker cell wall. The present study showed geographical variation with regards to MLST profiles, antimicrobial resistance, and virulent factors in global *M. caseolyticus* lineages. This study suggests that such local evolution of foodborne or livestock origin *M. caseolyticus* may serve as vehicles for domestic transmission of methicillin resistance in retail meat in China.

**Highlights:**

- Global *M. caseolyticus* strains were divided into five lineages from A to E.
- MLST typing revealed novel alleles in Chinese *M. caseolyticus* strains.
- Chinese lineage carried significantly more AMR genes and less virulent factors.
- *Macrococcus caseolyticus* has a larger diameter and thicker cell wall compared with *S*.*aureus*.
- *Macrococcus caseolyticus* may enhance the domestic transmission of methicillin resistance in China.

## Introduction

The genus *Macrococcus* is a catalase-positive cocci that was first isolated in 1916 and currently consists of 12 species, including the classic *Macrococcus caseolyticus, Macrococcus bovicus, Macrococcus canis*, and other novel species (Evans, 1916; Mašlaňová et al., 2018). *Macrococcus caseolyticus*, the closest relative of staphylococci, contained a much smaller chromosome (2.1 MB) and lacks several sugar and amino acid metabolic pathways, as well as specific virulent genes that are commonly present in *S. aureus* (Mannerová et al., 2003; Baba et al., 2009).

*Macrococcus caseolyticus* is an adventitious bacterium that was frequently detected in European fermented cheeses and sausages, and enhanced the flavor of dairy products by producing amino acid and lipid-derived flavor compounds (Rantsiou et al., 2005; Fuka et al., 2013; Joishy et al., 2019). However, researches over the past decades have observed methicillin resistances *M. caseolyticus* with adaptive evolution of genes, *i*.*e*., *mecA, mecB, mecC*, and *mecD* (Ubukata et al., 1989; Tsubakishita et al., 2010; García-Álvarez et al., 2011; Schwendener et al., 2017), following a similar mechanism as MRSA. The classical *mecA* and *mecC* are generally considered to be present in *S. aureus* while homologous complexes *mecB* and *mecD* have been described in *M. caseolyticus* in 2012 and 2017, respectively. The *mec* complex was associated with transposon Tn6045 on the chromosome of *M. caseolyticus* JCSC7096, revealing a possibility of horizontal genetic transfers within the *Macrococcus* genus (Gómez-Sanz et al., 2015).

Recent studies also reported cases of macrococci associated with high mortality rates in veterinary, indicating the potential pathogenicity of infection in animals (Brawand et al., 2017; Cotting et al., 2017; Li et al., 2018). For instance, the *M. caseolyticus* SDLY strain isolated from commercial chickens extends severe pathogenicity including hemorrhages and multifocal necrosis, which was claimed to be associated with the insertion of capsular polysaccharide genes in a virulence context (Li et al., 2018).

To characterize the foodborne or livestock origin of methicillin-resistant *M. caseolyticus* from retail meat sold in Shanghai, China, we isolated and genomic sequenced 10 isolates from Shanghai, and compared them with 87 additional global isolates that were publicly available from the GenBank. Furthermore, we examined the phenotypic antimicrobial resistance profiles of Chinese isolates and observed morphological differences between *M. caseolyticus* and *S. aureus* to better understand these two bacteria species.

## Method

### Sampling and collection

Samples from each of 24 outlets, including 11 in wet markets and 13 in the supermarkets, were collected between August and October of 2021 in Shanghai (Figure 1). The sampled outlets were distributed across 11 districts from the Shanghai city, including Pudong (n=6), Huangpu (n=3), Xuhui (n=2), Jing’an (n=2), Minhang (n=2), Yangpu (n=2), Changning (n=2), Putuo (n=2), Songjiang (n=1), Fengxian (n=1), and Baoshan (n=1). Packages of frozen chicken, beef, and pork were purchased from the outlets and transported to the laboratory on ice containers within 4 h of collection. Retail meat products were prepared for analysis as previously described (Li et al., 2021). Briefly, 10 g of samples were homogenized in 0.1% peptone saline in a filter bag (Bkmam, Changde, China) and 100 ul were cultured onto CHROMagar™ MRSA agar (Becton Dickinson, Franklin Lakes, NJ) for selection overnight at 37°C (Keller et al., 2022). The obtained *M. caseolyticus* strains were confirmed by 16s rRNA sequencing and MALDI-TOF MS (Bio-M erieux, Craponne, France).

**Figure 1.**
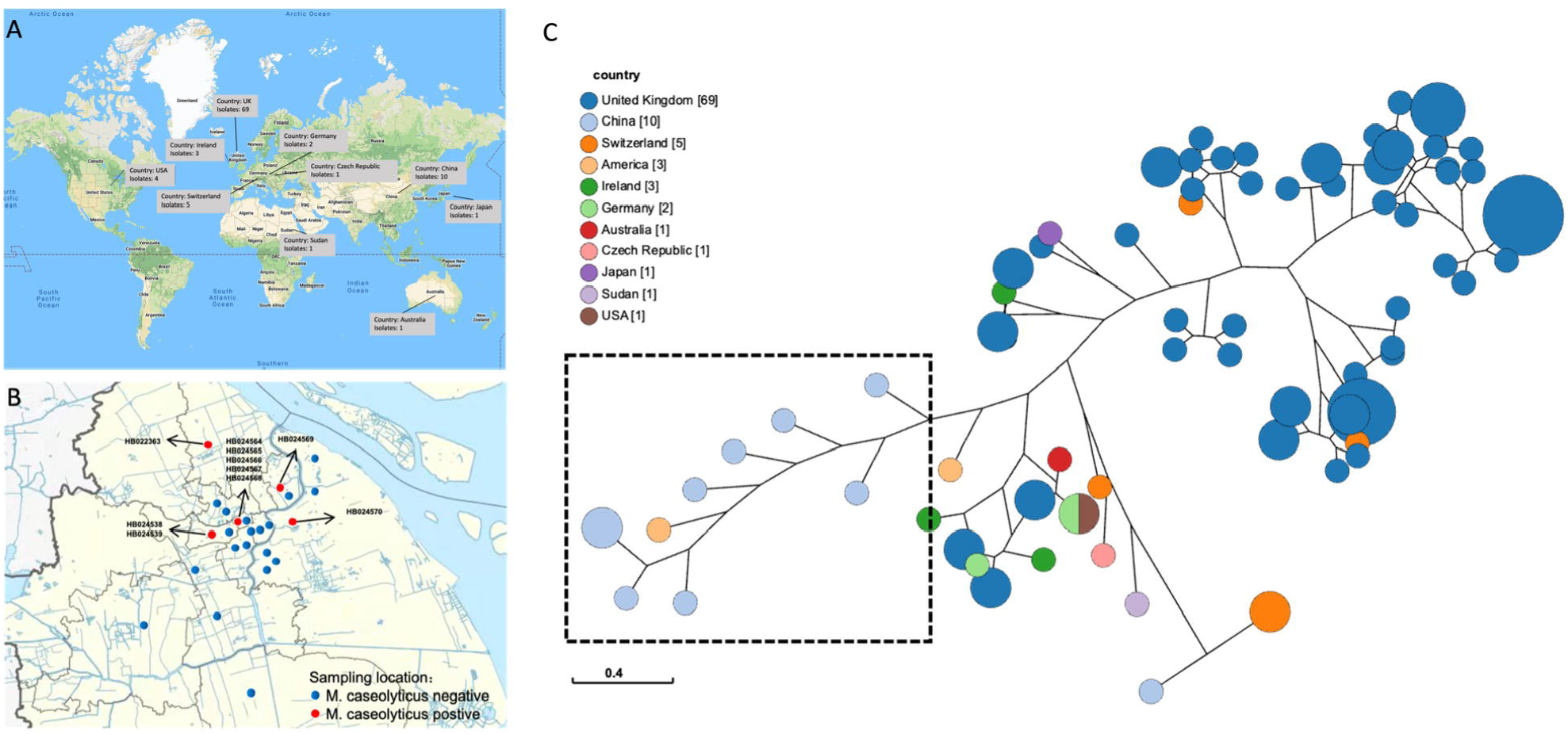
Geographic distribution and phylogenetic analysis of the *M. caseolyticus*. (A) Global distribution of 97 *M. caseolyticus* isolates. (B) Geographical distribution of 10 Chinese *M. caseolyticus* isolates in this study. (C) Phylogenetic analysis of 97 *M. caseolyticus* isolates in the global lineage. The black dashed box indicated the Chinese lineage except for HB022363 isolates.

### Antimicrobial susceptibility test

Confirmed *M. caseolyticus* strains were enrolled for antimicrobial susceptibility test by disk diffusion (Oxoid) according to the guideline of Clinical and Laboratory Standards Institute (CLSI 2017). The antimicrobial compounds included ampicillin (10 ug), amikacin (30 ug), cefazolin (30 ug), cefuroxime (30 ug), ceftriaxone (30 ug), ceftazidime (30 ug), cefoperazone (75 ug), doxycycline (30 ug), erythromycin (15 ug), gentamicin (10 ug), kanamycin (30 ug), lincomycin (2 ug), minocycline (30 ug), penicillin (10 ug), piperacillin (100 ug), streptomycin (10 ug), tetracycline (30 ug), and with cefoxitin as methicillin-resistant drug control.

### Whole genome sequencing

A total of 10 *M. caseolyticus* strains isolated from this study were sent for DNA purification using HiPure Bacterial DNA Kit (D3146, Meiji Biotechnology Co., Ltd, Guangzhou, China) and sequenced via Illumina’s NextSeq 500 (Illumina, San Diego, CA, United States) on the platform of Honsunbio company (Shanghai, China). In addition, 87 global strains were downloaded from GenBank (https://www.ncbi.nlm.nih.gov/genbank/), including 53 assemblies and 34 sets of raw reads. All the raw reads were trimmed and assembled by EToKi, and quality was evaluated with QUAST v2.3 (Gurevich et al., 2013; Zhou et al., 2020). This resulted to a global collection of genomes of 97 *M*. *caseolyticus* strains. The raw reads from this study are submitted to the China National GenBank under project accession ID CNP0002826.

### Bioinformatic analysis

All 97 *M*. *caseolyticus* assemblies were submitted for MLST typing (https://pubmlst.org), and screened for antimicrobial-resistant genes and virulent factors using online tools of ResFinder, MobileElementFinder, and VFDB (Liu et al., 2019; Bortolaia et al., 2020; Johansson et al., 2021). Alignments with a minimum of ≥60% nucleotide identity was kept in all three programs, respectively. The single nucleotide polymorphism (SNP) tree was constructed using CSI Phylogeny v1.4 (http://genomicepidemiology.org/) by aligning other genomes onto a reference genomic sequences from *Macrococcus* sp. (ASM2009713). The genotypic data were visualized in Grapetree and ITOL (Zhou et al., 2018; Letunic and Bork, 2019). The violin graph was drawn in GraphPad Prism 7 and statistical significance was calculated using One-way ANOVA with p < 0.005.

### Comparative electron microscope observation

Prior to scanning and transmission electron microscope (SEM/TEM), both *M. caseolyticus* and *S. aureus* cultures were grown in 50 ml of LB at 37C with a starting OD600 of 0.02. Cells were harvested at OD600 ∼ 0.5 and suspended in fixation solution and incubated overnight at 4°C. After the treatments, cell cultures were washed twice with cacodylate buffer (0.05 M, pH 7.4) and post-fixed with 2% osmium tetroxide, followed by 0.25% uranyl acetate for contrast enhancement. The pellets were dehydrated with ethanol (30, 50, 70, 80, 90, and 100%), embedded in Epoxy resin, and cut into ultrathin sections for lead citrate staining. The final sections were examined by Philips CM100 BioTWIN transmission electron microscope. For the SEM, the suspended cells after post-fixation and dehydration were placed on stubs and coated with gold-palladium for 2 min. Then samples were observed by JSM-7500F scanning electron microscope (JEOL Ltd., Tokyo, Japan).

## Results

### Geographic distribution and population structure

A total of 97 *M. caseolyticus* strains were selected for geographic analysis including 10 isolates from Shanghai and 87 isolates from GenBank (Figure 1). All strains were distributed in five continents including Europe (82.4%, n = 80), Asia (11.3%, n = 11), North America (4.1%, n = 4), Oceania (1%, n = 1) and Africa (1%, n = 1) (Figure 1). The phylogenetic tree showed that global *M. caseolyticus* strains were divided into five lineages from A to E. Lineage A forms an independent cluster that was separated from all the rest strains. Lineage B are the specific strains that associated with human infections, while lineage C from this study was mainly detected in retail meat (beef, n=6; pork, n=4) as a local Chinese cluster except for HB022363 isolates. Lineage D and E have the dominant proportion isolated from bulk milk (Figure 2). MLST typing of lineage B also showed that several novel alleles were identified among the Chinese *M. caseolyticus* strains, e.g., *fdh* and *purA* in HB022363 and *cpn60* and *pta* in HB024569 (Table 1 & Supp. Table 1).

**Table 1.** Multi-locus sequence typing (MLST) of 10 *M. caseolyticus* strains isolated from China.

| 456 | MLST | ack | cpn60 | fdh | pta | purA | sar | tuf |
| --- | --- | --- | --- | --- | --- | --- | --- | --- |
| 457 | HB022363 | 14 | 16 | 18* | 4 | 13*/15* | 17 | ND |
| 458 | HB024538 | 6 | 2 | 9 | 5* | 3 | 7 | 4 |
| 459 | HB024539 | 6 | 13* | 9* | 10* | 3* | 7 | 5 |
| 460 | HB024564 | 6 | 13* | 9* | 5 | 20* | 7 | 5 |
| 461 | HB024565 | 6 | 2 | 9* | 10* | 20* | 17* | 5 |
| 462 | HB024566 | 6 | 2*/12* | 9* | 5 | 20* | 1 | 5 |
| 463 | HB024567 | 6 | 2*/12* | 9* | 5 | 20* | 1* | 5 |
| 464 | HB024568 | 6 | 2* | 9* | 5 | 20* | 2* | 5 |
| 465 | HB024569 | 6 | 3* | 3 | 1* | 3 | 1 | 4 |
| 466 | HB024570 | 6 | 3* | 9* | 5 | 2 | 1 | 5 |
| 467 |  |  |  |  |  |  |  |  |

Asterisks indicate novel alleles with the closest types. ND: Not Detected.

**Figure 2.**
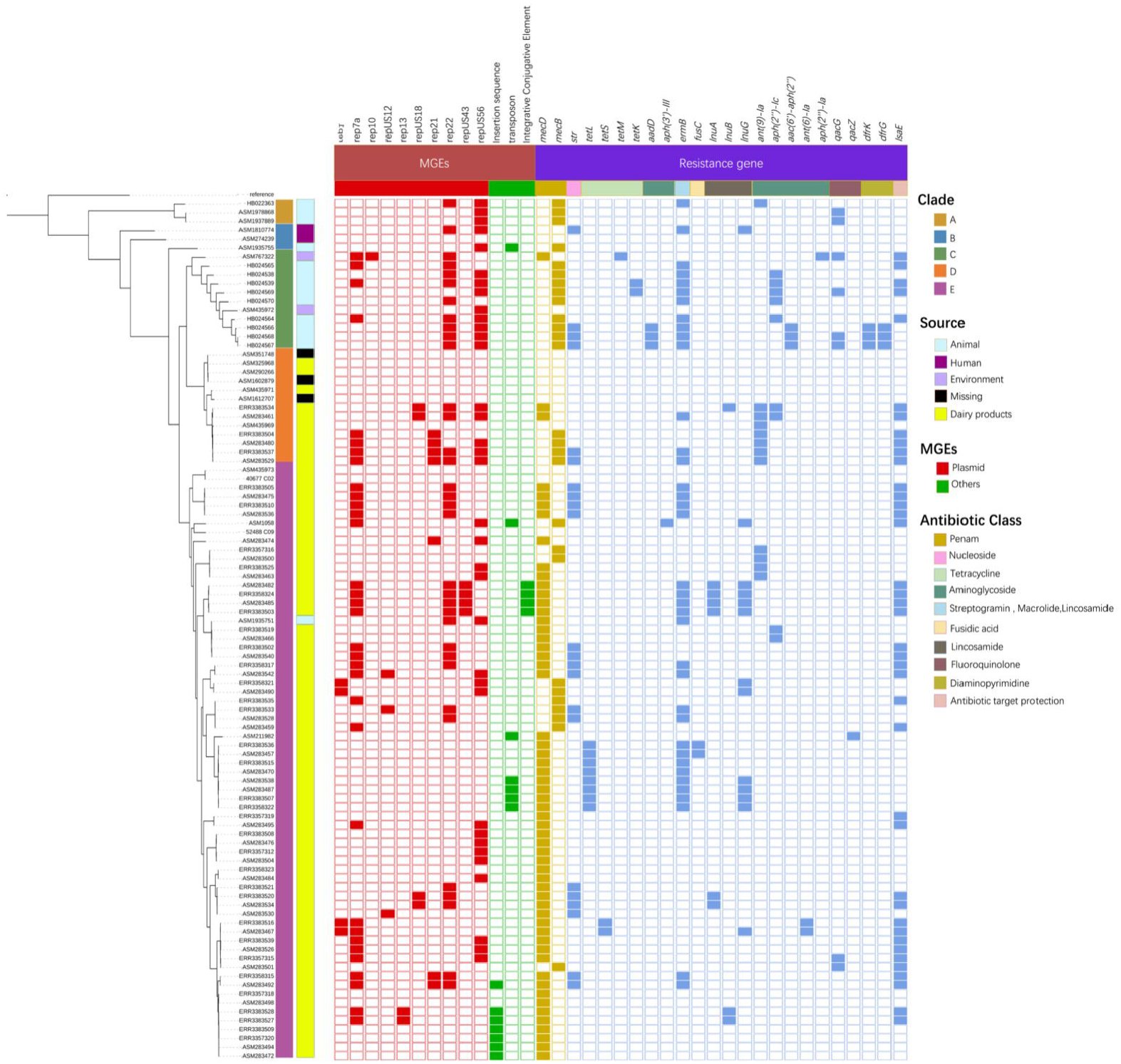
Distributions of MGEs and AMR genes in *M. caseolyticus* lineages. *Macrococcus caseolyticus* were divided into five lineages including A, B, C, D and E based on phylogenetic tree. The squares colored by trait category represented the presence of MGE or AMR genes.

### Identification of AMR genes and mobile elements

Given the importance of antimicrobial resistance (AMR) in opportunistic pathogens, we compared the distributions of AMR genes in 97 *M. caseolyticus* strains from different lineages (Figure 2). A total of 24 AMR genes associated with 10 classes of antimicrobial agents were present in the isolates from five lineages, among which 33% (1/3, lineage A), 33% (1/3, lineage B), 91% (10/11, lineage C), 46% (6/13, lineage D) and 43% (29/67, lineage D) of isolates from the corresponding lineages were identified as multi-drug resistant (Figure 2). Notably, all the 10 Chinese isolates contained *mecB* gene whiles HB024567, HB024568, and HB024569 carried several unique genes including *drfK, drfG, aph(3’)-III*, and *aac(6’)-aph(2’’)*. In addition, *tetk* was also detected uniquely in HB024539 and HB024569 which may be associated with the horizon plasmid transfer of repUS56. The mobile genetic elements analysis showed that rep7a, rep22, and repUS56 were the top dominant plasmids in *M. caseolyticus* with the prevalence of 34%, 35%, and 37% in global lineages (Figure 2). Further statics indicated that the number of AMR genes from lineage C was significantly higher than those from lineage D and E (p<0.005, Figure 3).

**Figure 3.**
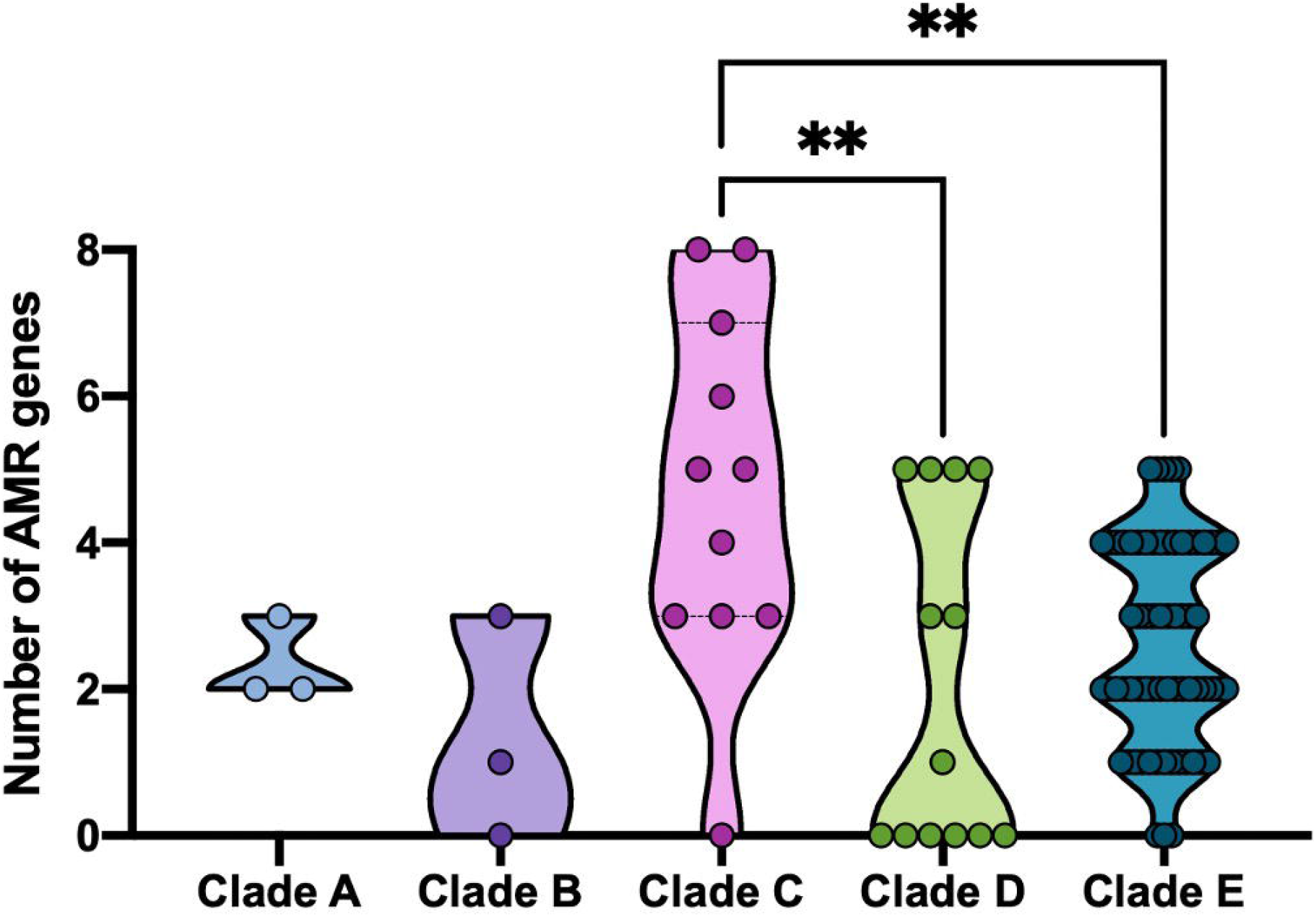
The number of AMR genes in *M. caseolyticus* lineages. **, P < 0.005.

### Phenotypical antimicrobial resistance of Chinese lineage

All 10 Chinese *M. caseolyticus* isolates were screened for phenotypic antimicrobial-resistant profiles. Specifically, all isolates were resistant to ampicillin, cefazolin, ceftazidime, lincomycin, piperacillin, penicillin, streptomycin, and tetracycline, whereas a majority were sensitive to amikacin, cefuroxime, and gentamicin, indicating a broad but complex multi-drug resistance in these *M. caseolyticus* strains isolated from retail meat in Shanghai, China (Table 2).

**Table 2.** Antimicrobial susceptibility of 10 *M. caseolyticus* isolates from China

| Antimicrobials | AMP | AMK | CZO | CXM | CRO | CAZ | CFP | DOXY | ERY | GEM | KAN | LIN | MNO | PEN | PIP | STR | TET |
| --- | --- | --- | --- | --- | --- | --- | --- | --- | --- | --- | --- | --- | --- | --- | --- | --- | --- |
| HB022363 | R | S | R | S | R | R | I | R | I | I | R | R | R | R | R | R | R |
| HB024538 | R | S | R | R | R | R | R | R | R | S | R | R | I | R | R | R | R |
| HB024539 | R | I | R | R | I | R | I | R | R | S | R | R | R | R | R | R | R |
| HB024564 | R | I | R | S | I | R | R | R | R | S | I | R | R | R | R | R | R |
| HB024565 | R | R | R | R | I | R | R | R | R | I | R | R | R | R | R | R | R |
| HB024566 | R | S | R | S | I | R | I | R | R | S | R | R | R | R | R | R | R |
| HB024567 | R | S | R | S | I | R | R | R | R | S | R | R | R | R | R | R | R |
| HB024568 | R | S | R | S | I | R | R | R | R | S | R | R | I | R | R | R | R |
| HB024569 | R | R | R | I | R | R | R | I | R | R | I | R | R | R | R | R | R |
| HB024570 | R | R | R | R | S | R | R | R | R | S | R | R | R | R | R | R | R |

S stands for sensitive, I for intermediate, and R for resistant.

### Distributions of virulent factors

Virulent factors were compared to determine the pathogenicity within the five lineages (Figure 4). Totally 79 virulent factors were assessed among 97 *M. caseolyticus* strains including the functional factors of adherence, biofilm formation, exotoxin, capsule, and others. Remarkably, the majority of lineage A and C isolates lack the factor of CTC01574 (hemolysin) but contained *hlyA* and CD1208 which are associated with hemolysin. Meanwhile, most lineage A and C strains contain cytolysin-related genes, e.g., *hpt, manA*, however, lack genes regulating capsule synthesis such as *capE, capF, capO*, and *capM* (capsular polysaccharide synthesis enzyme), indicating the weak ability of invasiveness. In lineage C, the replacement and loss of functional genes were also observed that KPHS_39850 (protein disaggregation chaperone) was only present in ASM767322 while A225_4443 (*clpB* factor) were widely distributed in Chinese isolates. Notably, the *aur* and *panD* genes were absent in Chinese isolates in lineage C. To summarize, the number of virulent factors in lineage C was significantly lower than that of lineage D and E (p<0.001), which may be due to the deletion of corresponding proteins related to capsule synthesis, zinc metalloproteinase aureolysin, and pantothenic acid synthesis (Figure 5).

**Figure 4.**
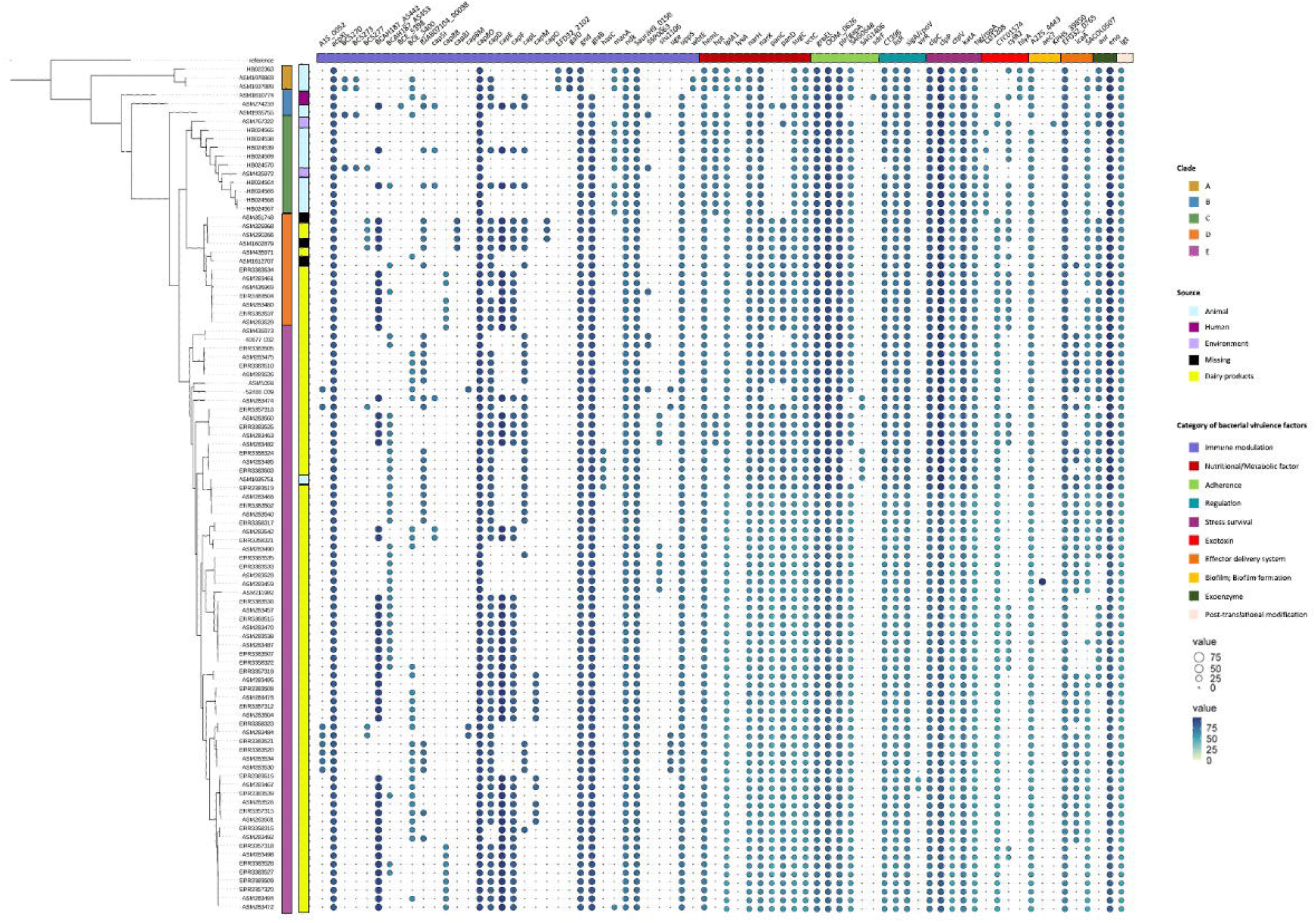
Distribution of virulent factors in the *M. caseolyticus* lineages. The larger the diameter of the circle and the darker the color, the higher the confidence value. Squares are colored by feature class, representing the presence of the checked features.

**Figure 5.**
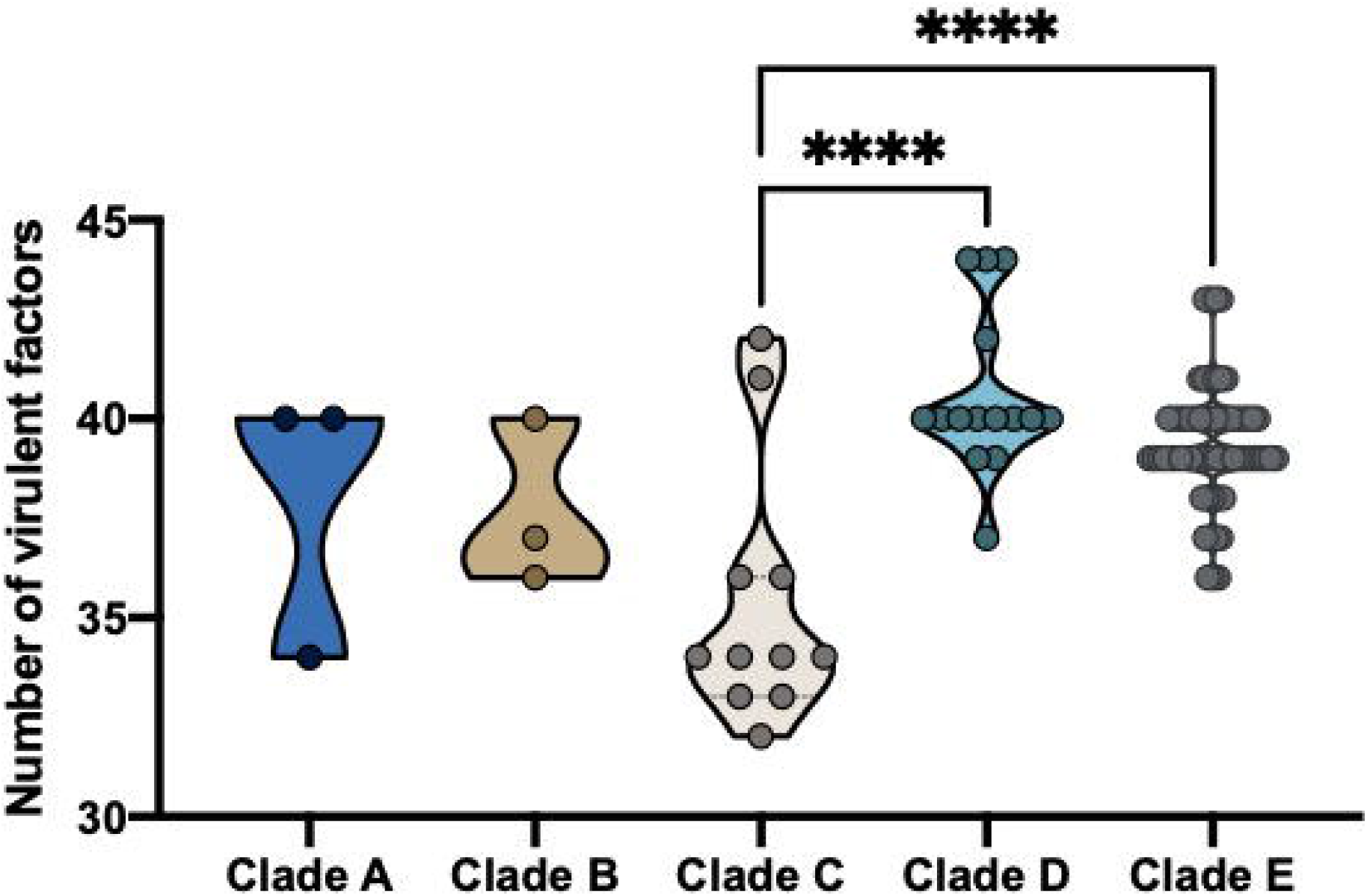
The number of virulent factors in *M. caseolyticus* lineages. ****, P < 0.001.

### Comparative morphological observation of *M. caseolyticus* and *S. aureus*

To better understand the morphological differences between *M. caseolyticus* and its closest relative of *S. aureus*, the HB024567 strain from present study and *S. aureus* ATCC 25923 were employed for SEM and TEM analysis. SEM results showed that *M. caseolyticus* was 1.1± 0.05 um with a round shape and smooth surface, while *S. aureus* was 0.46 ± 0.01 um which was smaller than the *M. caseolyticus* strain (Figure 6). Then the TEM results demonstrated that *M. caseolyticus* had a broad and thick cell wall with a diameter of around 65 ± 5 nm, whereas the cell wall was much narrow and thin in *S. aureus* (21 ± 1 nm) (Figure 7).

**Figure 6.**
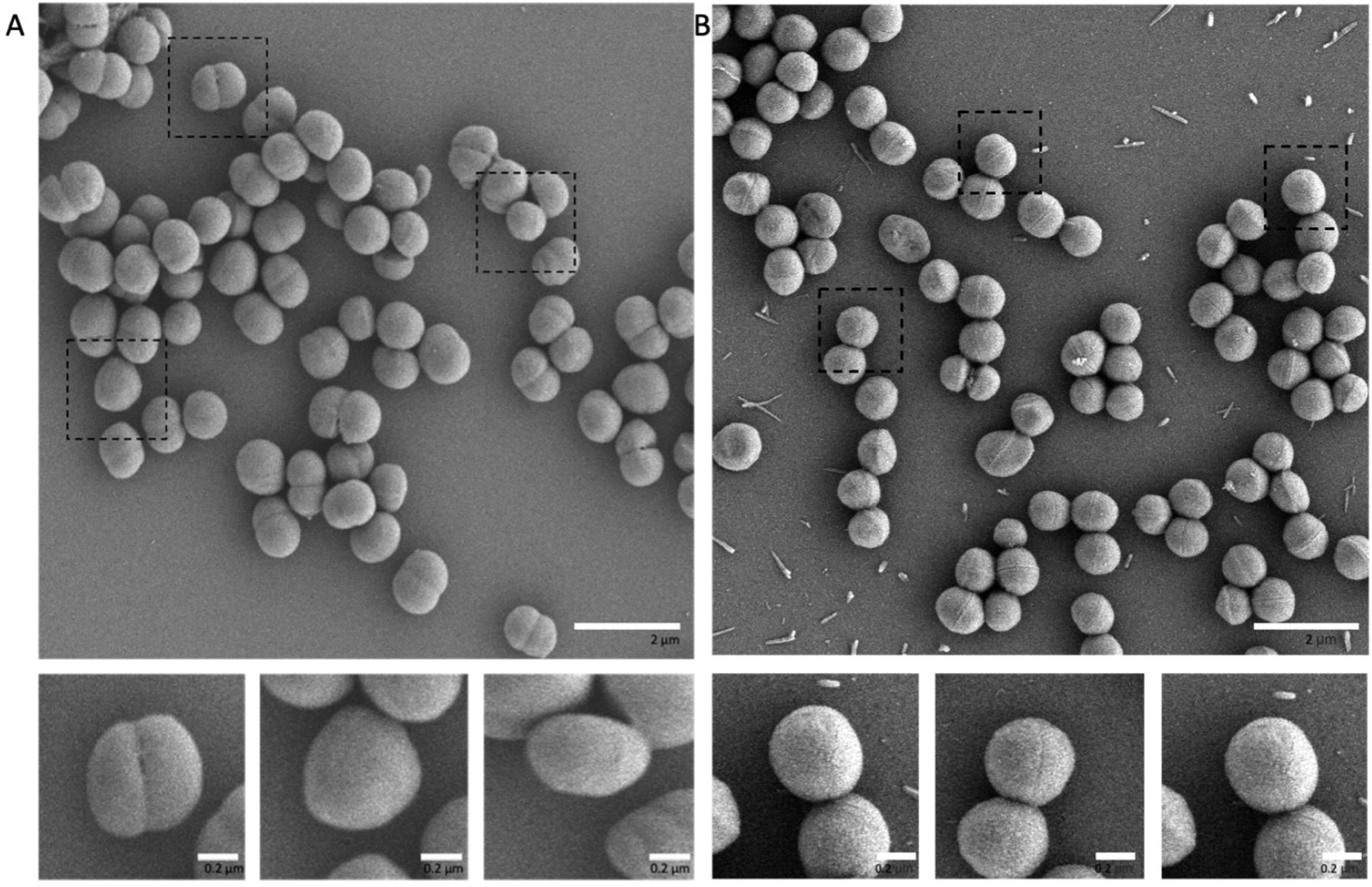
Scanning electron micrographs of (A) *M. caseolyticus* and (B) *S*.*aureus* strain.

**Figure 7.**
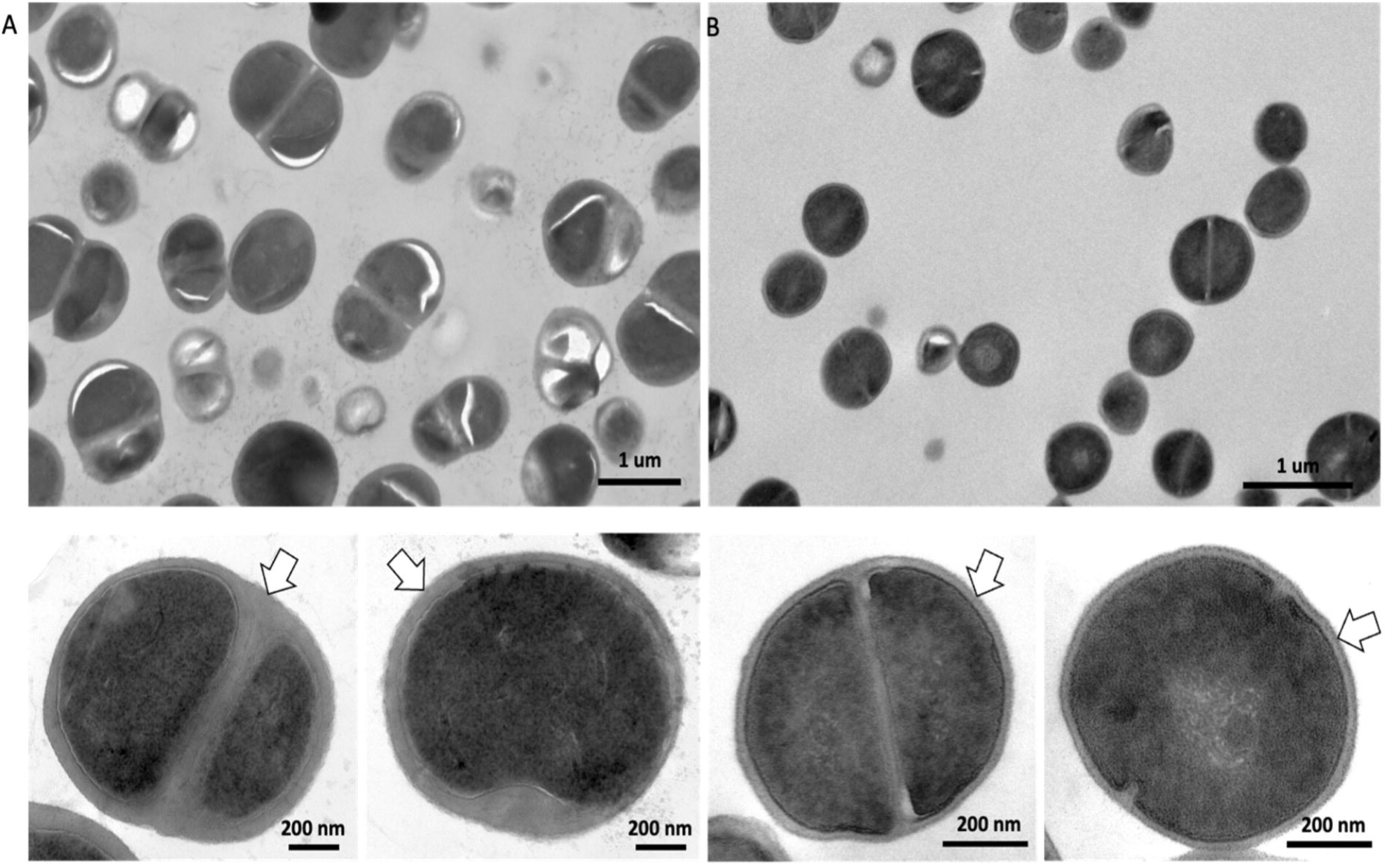
Transmission electron micrographs of (A) *M. caseolyticus* and (B) *S*.*aureus* strain.

## Discussion

*Macrococcus caseolyticus* is the foodborne bacterium detected in several fermented foods in China such as the Chinese sausages and the specialty food “ChouGuiYu”, a type of fermented mandarin fish (Dai et al., 2013). Previous studies also described the colonization of *M. caseolyticus* on the surface of humans, animals, and raw meat during food processing and transportation (Acheampong et al., 2021; Jost et al., 2021; Keller et al., 2022), indicating that trade in agricultural products may lead to the bacterial transfer across countries.

In the present study, *M. caseolyticus* were detected in raw meat with 8 isolates from beef, and the remaining 2 strains were from pork. Nine isolates of *M. caseolyticus* were distributed in lineage C as a cluster. According to the database of pubMLST, all 10 Chinese isolates showed novel alleles, and six of them contained more than three novel alleles. Taken together with the results of phylogenetic tree, such population structure may be due to the local evolution of *M. caseolyticus* in China. Notably, the HB022363 strain was separated from other Chinese isolates but clustered with ASM1978868 and ASM1937889 that were isolated from nasal swabs of the calf in Switzerland. Although the genomic analysis indicated that all three isolates in lineage A were closest relative to *M. caseolyticus*, further studies are needed to confirm their species and corresponding microbiological traits to explain the independent lineage in the phylogenetic tree.

*Macrococcus caseolyticus* is methicillin-resistant based on the presence of *mecB* and *mecD* elements. As the homologs of *mecA* gene, *mecB* and *mecD* were initially identified in *M. caseolyticus* strains in 2012 and 2017 (Schwendener et al., 2017). However, a recent study also reported the presence of *mecB* in *S. aureus* strains, indicating the potential horizon element transfer between these two species (Becker et al., 2018). In the present study, all 10 Chinese isolates were *mecB* positive with six of them identified as multidrug-resistant strains. Other AMR genes were evidenced to be carried by mobile genetic elements such as *str* gene in rep7a, *aadD*, and *tetL* genes in rep22. Intriguingly, lineage C had the highest number of AMR genes compared with other lineages, which revealed an increasing antimicrobial resistance in retail raw meat in Shanghai, China.

In addition to antimicrobial-resistant genes, the characteristics of virulent factors were screened for global lineages. The Chinese *M. caseolyticus* isolates were clustered together with ASM767322 and ASM435972. Comparative genomic analysis indicated the replacement and loss of several functional genes between ASM767322 and the rest isolates from lineage C. Notably, genes related to capsule synthesis were absent in Chinese isolates, which were previously reported as the dominant virulent factors in broiler disease (Li et al., 2018). The *panC* and *panD* genes that functioned in pantothenic acid synthesis also vanished in the present Chinese isolates. In summary, the decrease of virulent factors in Chinese isolates indicated weaker pathogenicity, and this may be due to the isolated environment of retail raw meat instead of human skin or nasal samples.

Scanning electron microscopy (SEM) and transmission electron microscopy (TEM) results showed that *M. caseolyticus* possessed a diameter of 1.1± 0.05 μm with a cell wall of 65 ± 5 nm (Figure 6 A & B), which showed a larger diameter and cell wall compared to *S. aureus*. The genus *Macrococcus* was measured in previous studies with the diameters of *M*.*caseolyticus* ATCC13548 (1.1-2 μm), *M*.*equipercicus* ATCC51831 (1.3-2.3 μm), *M. bovicus* ATCC51825 (1.2-2.1 μm), *M*.*carouselicus* ATCC5128 (1.4-2.5 μm), *M. brunensis* CCM4811 (0.89 -1.2 μm), *M. hajekii* CCM4809 (0.89 μm), *M. lamae* CCM4815 (0.74 -0.92 μm), and *M. canis* KM45013 (0.8 μm) (Evans, 1916; Kloos et al., 1998; Mannerová et al., 2003; Brawand et al., 2017). SEM results showed that *M. caseolyticus* had a smooth surface, while it was rough in *M. bovicus, M. equipercicus*, and *M. carouselicus*, with spiny protrusions observed in *M. equipercicus* stains (Kloos et al., 1998).

In conclusion, comparative genomic analysis was enrolled for the construction of population structure, and prediction of antimicrobial resistance and virulent factors in 97 *M. caseolyticus* strains collected from humans, animals, meat, and dairy products. Five lineages were identified globally that Chinese isolates were clustered together as lineage C that carried significantly more AMR genes (p<0.005) and less virulent factors (p<0.001). MLST typing and phylogenetic tree indicated a potential local evolution of *M. caseolyticus* in China. The comparative electron microscope demonstrated the morphological changes between *M. caseolyticus* and *S. aureus*, showing that *M. caseolyticus* has a larger diameter and thicker cell wall. Together, the present study provided a phylogenetical and genotypical comparison for global *M. caseolyticus* stains. Our results show that both human and animal reservoirs can contribute to contamination in food products and that trade in agricultural products may serve as a vehicle for *M. caseolyticus* transfer in China.

