## Supplementary material for "Genomic analysis of population structure, antimicrobial resistance, and virulent factors of methicillin-resistant *Macrococcus caseolyticus* in global lineages": Supp. Table 1

Supply table 1. Multi-locus sequence typing (MLST) identified in 87 global *M. caseolyticus* isolates.

| MLST ST | ack | cpn60 | fdh | pta | purA | sar | tuf |
| --- | --- | --- | --- | --- | --- | --- | --- |
| ERR3357312 | 6 | 11 | 11 | 2 | 12 | 3 | 2 |
| ERR3357315 | 6 | 4 | 11 | 2 | 5 | 3 | 2 |
| ERR3357316 | 15 | 3 | 11 | 2 | 5 | 3 | 3 |
| ERR3357318 | 6 | 4 | 5 | 2 | 5 | 3 | 2 |
| ERR3357319 | 15 | 13 | 13 | 2 | 12 | 6 | 2 |
| ERR3357320 | 6 | 4 | 5 | 2 | 5 | 3 | 2 |
| ERR3358315 | 6 | 4 | 15 | 2 | 5 | 3 | 3 |
| ERR3358317 | 6 | 3 | 11 | 2 | 5 | 3 | 3 |
| ERR3358321 | 6 | 3 | 5 | 2 | 5 | 6 | 3 |
| ERR3358322 | 5 | 3 | 5 | 2 | 5 | 3 | 2 |
| ERR3358323 | 4 | 4 | 5 | 2 | 5 | 3 | 2 |
| ERR3358324 | 5 | 9 | 5 | 7 | 10 | 6 | 2 |
| ERR3383502 | 6 | 3 | 5 | 7 | 5 | 6 | 2 |
| ERR3383503 | 5 | 9 | 5 | 7 | 10 | 6 | 2 |
| ERR3383504 | 6 | 3 | 4 | 3 | 6 | 3 | 2 |
| ERR3383505 | 6 | 8 | 5 | 7 | 5 | 14 | 2 |
| ERR3383507 | 5 | 3 | 5 | 2 | 5 | 3 | 2 |
| ERR3383508 | 6 | 11 | 11 | 2 | 12 | 3 | 2 |
| ERR3383509 | 6 | 4 | 5 | 2 | 5 | 3 | 2 |
| ERR3383510 | 6 | 8 | 5 | 7 | 5 | 14 | 2 |
| 40677_C02 | 11 | 3 | 5 | 7 | 8 | 3 | 3 |
| 52488_C09 | 6 | 6 | 8 | 2 | 8 | 1 | ND |
| ASM1058v1 | 3 | 3 | 15*/8* | 7 | 3 | 11 | 2 |
| ASM211982v1 | 5 | 3 | 5 | 2 | 5 | 3 | 2 |
| ASM274239v2 | 13* | 13* | 7* | 4* | 2/14 | 17* | 12 |
| ASM283457v1 | 5 | 3 | 5 | 2 | 5 | 3 | 2 |
| ERR3383515 | 5 | 3 | 5 | 2 | 5 | 3 | 2 |
| ERR3383516 | 6 | 4 | 11 | 2 | 5 | 3 | 9 |
| ERR3383519 | 6 | 3 | 5 | 7 | 5 | 6 | 2 |
| ERR3383520 | 4 | 4 | 5 | 2 | 5 | 3 | 2 |
| ERR3383521 | 4 | 4 | 5 | 2 | 5 | 3 | 2 |
| ERR3383525 | 15 | 3 | 11 | 2 | 5 | 3 | 3 |
| ERR3383527 | 6 | 4 | 5 | 2 | 5 | 3 | 2 |
| ERR3383528 | 6 | 4 | 5 | 2 | 5 | 3 | 2 |
| ERR3383533 | 6 | 4 | 5 | 2 | 5 | 3 | 2 |
| ERR3383534 | 6 | 3 | 4 | 3 | 6 | 3 | 2 |
| ERR3383535 | 16 | 3 | 5 | 7 | 17 | 6 | 2 |
| ERR3383536 | 5 | 3 | 5 | 2 | 5 | 3 | 2 |
| ERR3383537 | 6 | 3 | 4 | 3 | 6 | 3 | 2 |
| ERR3383539 | 6 | 4 | 11 | 2 | 5 | 3 | 2 |
| ASM283459v1 | 16 | 3 | 5 | 7 | 17 | 6 | 2 |
| ASM283461v1 | 6 | 3 | 4 | 3 | 6 | 3 | 2 |
| ASM283463v1 | 15 | 3 | 11 | 2 | 5 | 3 | 3 |
| ASM283466v1 | 6 | 3 | 5 | 7 | 5 | 6 | 2 |
| ASM283467v1 | 6 | 4 | 11 | 2 | 5 | 3 | 9 |
| ASM283470v1 | 5 | 3 | 5 | 2 | 5 | 3 | 2 |
| ASM283472v1 | 6 | 4 | 5 | 2 | 5 | 3 | 2 |
| ASM283474v1 | 16 | 3 | 16 | 2 | 16 | 3 | 2 |
| ASM283475v1 | 6 | 8 | 5 | 7 | 5 | 14 | 2 |
| ASM283476v1 | 6 | 11 | 11 | 2 | 12 | 3 | 2 |
| ASM283480v1 | 6 | 3 | 4 | 3 | 6 | 3 | 2 |
| ASM283482v1 | 5 | 9 | 5 | 7 | 10 | 6 | 2 |
| ASM283484v1 | 4 | 4 | 5 | 2 | 5 | 3 | 2 |
| ASM283485v1 | 5 | 9 | 5 | 7 | 10 | 6 | 2 |
| ASM283487v1 | 5 | 3 | 5 | 2 | 5 | 3 | 2 |
| ASM283490v1 | 6 | 3 | 5 | 2 | 5 | 6 | 3 |
| ASM283492v1 | 6 | 4 | 15 | 2 | 5 | 3 | 3 |
| ASM283494v1 | 6 | 4 | 5 | 2 | 5 | 3 | 2 |
| ASM283495v1 | 15 | 13 | 13 | 2 | 12 | 6 | 2 |
| ASM283498v1 | 6 | 4 | 5 | 2 | 5 | 3 | 2 |
| ASM283504v1 | 6 | 11 | 11 | 2 | 12 | 3 | 2 |
| ASM283526v1 | 6 | 4 | 11 | 2 | 5 | 3 | 2 |
| ASM283528v1 | 16 | 3 | 5 | 7 | 17 | 6 | 2 |
| ASM283529v1 | 6 | 3 | 4 | 3 | 6 | 3 | 2 |
| ASM283530v1 | 4 | 4 | 5 | 2 | 5 | 3 | 2 |
| ASM283534v1 | 4 | 4 | 5 | 2 | 5 | 3 | 2 |
| ASM283536v1 | 6 | 8 | 5 | 7 | 5 | 14 | 2 |
| ASM283538v1 | 5 | 3 | 5 | 2 | 5 | 3 | 2 |
| ASM283540v1 | 6 | 3 | 5 | 7 | 5 | 6 | 2 |
| ASM283542v1 | 6 | 3 | 11 | 2 | 5 | 3 | 3 |
| ASM290266v1 | 3 | 3 | 4 | 3 | 4 | 3 | 3 |
| ASM325968v1 | 3 | 3 | 4 | 3 | 4 | 3 | 3 |
| ASM351748v1 | 3 | 3 | 4 | 3 | 4 | 17* | 3 |
| ASM435969v1 | 6 | 3 | 4 | 3* | 6 | 3 | 2 |
| ASM435971v1 | 6 | 3 | 6 | 3 | 7 | 3 | 3 |
| ASM435972v1 | 8 | 3 | 9* | 5 | 2 | 1 | 5 |
| ASM435973v1 | 11 | 3 | 5 | 7 | 8 | 3 | 3 |
| ASM1602879v1 | 3 | 3 | 4 | 3 | 4 | 3 | 3 |
| ASM1612707v1 | 6 | 3 | 5 | 3 | 7 | 3 | 3 |
| ASM1810774v1 | 6 | 17* | 10 | 3* | 2* | 17 | 12 |
| ASM1935751v1 | 6 | 3 | 5 | 7 | 5 | 6 | 2 |
| ASM1935755v1 | 6 | 6 | 7 | 6 | 10 | 5 | 6 |
| ASM1937889v1 | 14 | 17 | 15 | 4 | 19 | 17 | 8 |
| ASM1978868v1 | 14 | 17 | 15 | 4 | 19 | 17 | 8 |
| ASM767322v1 | 13 | 1 | 2 | 1 | 1 | 1 | 1 |
| ASM283500v1 | 15 | 3 | 11 | 2 | 5 | 3 | 3 |
| ASM283501v1 | 6 | 4 | 11 | 2 | 5 | 3 | 2 |

Asterisks indicate novel alleles with the closest types. ND: Not Detected.
